# Single-cell RNA-seq reveals trans-sialidase-like superfamily gene expression heterogeneity in *Trypanosoma cruzi* populations

**DOI:** 10.1101/2025.01.14.633000

**Authors:** Lucas Inchausti, Lucía Bilbao, Vanina A. Campo, Joaquín Garat, José Sotelo-Silveira, Gabriel Rinaldi, Virginia M. Howick, María Ana Duhagon, Javier G. De Gaudenzi, Pablo Smircich

## Abstract

*Trypanosoma cruzi*, the causative agent of Chagas disease, presents a major public health challenge in Central and South America, affecting approximately 8 million people and placing millions more at risk. The *T. cruzi* life cycle includes transitions between epimastigote, metacyclic trypomastigote, amastigote, and blood trypomastigote stages, each marked by distinct morphological and molecular adaptations to different hosts and environments. Unlike other trypanosomatids such as *Trypanosoma brucei*, *T. cruzi* does not employ a monoallelic model of antigenic variation; instead, it relies on a diverse repertoire of cell-surface associated proteins encoded by large multigene families, which are essential for infectivity and immune evasion.

This study analyzes cell-specific transcriptomes using single-cell RNA sequencing of amastigote and trypomastigote cells to characterize stage-specific surface protein expression during mammalian infection. Through clustering and identification of cell-specific markers, we assigned cells to distinct parasite developmental forms. Analysis of individual cells revealed that surface protein-coding genes, especially members of the trans-sialidase like superfamily (TcS), are expressed with greater heterogeneity than single-copy genes. Moreover, no recurrent combinations of TcS genes were observed between individual cells in the population. Remarkably, a small subset of TcS mRNAs, encoded by genes preferentially located in the core genomic compartment, are frequently detected across the cell population, whereas the vast majority of TcS mRNAs show low detection frequencies and are mainly encoded in the disruptive compartment. Our findings thus reveal transcriptomic heterogeneity within trypomastigote populations where each cell displays unique TcS expression profiles. Focusing on the diversity of surface protein expression, this research aims to deepen our understanding of *T. cruzi* cellular biology and infection strategies.

## Introduction

The protozoan parasite *Trypanosoma cruzi* is the etiological agent of Chagas disease, a highly prevalent infectious disease in Central and South America that affects approximately 8 million people, with several million more at risk of infection (1).

The parasite life cycle involves both an invertebrate vector (triatomine bug) and a vertebrate host. The change in environmental conditions triggers differentiation processes in the parasite developing across four main stages. The epimastigote form replicates within the insect and differentiates into metacyclic trypomastigotes in the insect’s rectal tract. This latter form does not replicate, specializes in host infection, and can invade various cell types. Once inside the cell, the parasite differentiates into an amastigote, the replicative form within the mammalian host. After several rounds of division, amastigotes differentiate into bloodstream trypomastigotes, which, after cell lysis, can either infect new cells or be ingested by the vector (2). Parasite developmental stages exhibit significant morphological and molecular differences, associated with distinct gene expression profiles. In trypanosomatids, protein-coding genes are organized into large polycistronic RNA polymerase II transcription units, with limited evidence for tight, gene-specific transcriptional control (3). As a result, gene expression is largely regulated at the post-transcriptional level, through mechanisms controlling mRNA steady-state abundance, translation, and related processes (3). More recently, accumulating evidence indicates that genome organization and chromatin state also shape transcriptional outputs (4–8). These studies have identified two distinct chromatin compartments in *T. cruzi*: a core compartment, in which genes tend to show more uniform expression consistent with permissive transcription and predominant post-transcriptional regulation (9), and a disruptive compartment, enriched in multigene families, whose genes exhibit lower average transcription and stronger chromatin-associated regulation (4).

*T. cruzi* infection relies on a heterogeneous set of membrane proteins, encoded mainly by large multigene families (10). Among these are trans-sialidases and trans-sialidase-like proteins (TcS), mucins, MASPs, GP63, DGFs, and RHS proteins, most of which are involved in infection, tropism, and immune evasion (11–16).

The trans-sialidase-like superfamily is involved in processes underlying host-parasite interactions (12). Members that retain enzymatic activity catalyze the transfer of sialyl groups from host glycoconjugates to galactopyranosyl units on the parasite’s surface, an essential activity, as *T. cruzi* is unable to synthesize *de novo* sialic acid (17). This is the largest superfamily, with over 1,400 members, and is subdivided into eight groups based on amino acid sequence (12). Few members have been functionally characterized, most expressed primarily in mammalian stages (18). The TcS group I include proteins that retain trans-sialidase activity, though members from all groups are involved in host-parasite interactions (19).

In recent years, single-cell RNA sequencing (scRNA-seq) has been employed in protozoa, with reports including *Plasmodium falciparum*, *Trypanosoma brucei,* and *Leishmania major* (20–29). These studies revealed key aspects of the infection process undetectable with conventional methods, highlighting the relevance of this approach for understanding individual variation in gene expression in single-celled organisms (30). In *T. brucei,* antigenic variation driven by variant surface glycoproteins (VSGs) has been studied at single-cell resolution to understand the mechanisms that enable subpopulations of this parasite to evade the immune system, as a bet hedging strategy, ensuring parasite survival (25). In this parasite, scRNA-seq revealed that pre-metacyclic cells express multiple VSG transcripts simultaneously, in contrast to metacyclic forms, which display a protein coat composed of a single VSG type.

While population-level heterogeneity in surface protein expression has been suggested as critical for *T. cruzi* infection, immune evasion, and persistence (31,32), this has not been studied at the intra-population level, and the underlying mechanisms remain poorly understood. Here, we present a scRNA-seq analysis of *T. cruzi*, to understand the heterogeneity in surface protein expression within trypomastigote populations.

## Results and Discussion

### Identification of cell populations

To assess the expression of cell-surface protein-coding genes in *Trypanosoma cruzi*, we conducted a 10X Chromium Single Cell 3’ assay from a mixed population of amastigotes and cell-derived trypomastigotes, aiming at sequencing the transcriptome of 5000 cells. After low-quality cell filtering and gene expression quantification (see Materials and Methods), we obtained 3192 single-cell transcriptomes with 14321 total genes detected, with a mean of 1088 genes and 2461 UMIs detected per cell. These results were comparable to other scRNA-seq studies done in *Trypanosoma brucei* and more recently addressed in *T. cruzi* (currently reported as a preprint) using 10X Chromium technology (26,33,34). Cell populations (**Figure 1a**) were defined by identifying cluster-specific gene markers (**Figure 1b and Supplementary File 1**). Two cell clusters were assigned to trypomastigotes and amastigotes: cluster 0 (2201 cells) and cluster 1 (824 cells), respectively. Markers gene expression was consistent with previous bulk RNA-seq data from Dm28c (4) (**Figures 1b, 1c and 1d, Supplementary Table 1**). In turn, we hypothesize that cluster 2, that comprised only 167 cells, reflects amastigote-trypomastigote transitioning parasites, as its gene markers are differentially expressed in bulk RNA-seq data, but some are upregulated in amastigotes and others in trypomastigotes (Supplementary Table 1).

**Figure 1.**
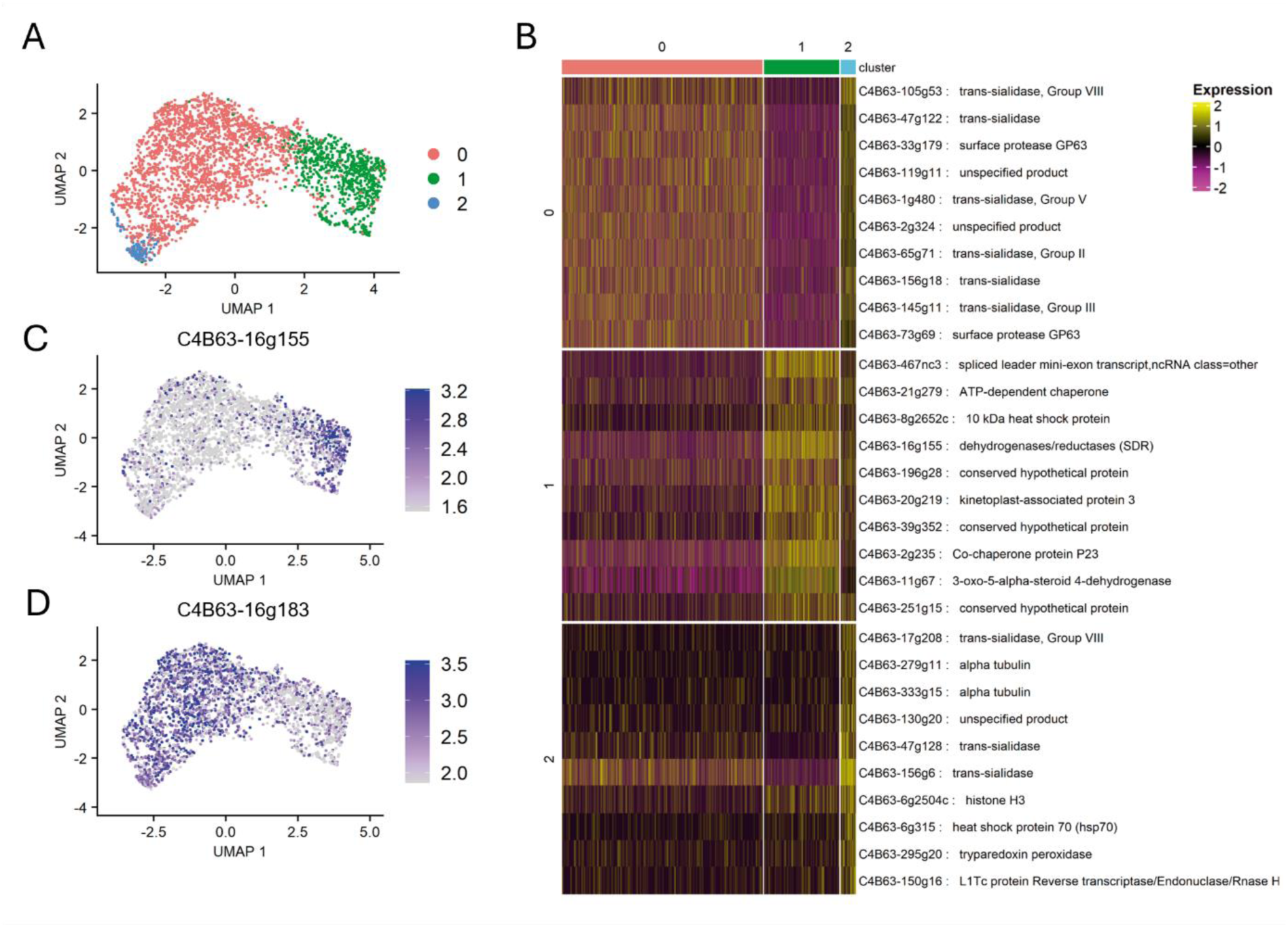
Identification of amastigote and trypomastigote cell populations. (a) UMAP colored by detected clusters based on gene expression profiles. (b) Heatmap of the top 10 gene markers upregulated in each of the 3 cell populations identified (log2FC > 1 and adjusted p-value < 0.05). (c) Expression of a cluster 0 marker gene (C4B63-16g183) on the UMAP, and (d) Expression of a cluster 1 marker gene (C4B63-16g155) on the UMAP.

### Expression pattern of surface protein-coding genes

We analyzed differences in gene expression between single-copy genes and multigene families, between and within the identified cell populations. As previously reported (4,18), multi gene family expression is increased in trypomastigotes (**Figure 2a**), consistent with the involvement of these genes in stage-specific functions. As control, we examined the expression of single-copy genes, for which this pattern was not observed as these genes are primarily associated with basic cellular functions and show more similar average expression levels between the two cell types (**Supplementary Figure 1a**)(33).

**Figure 2.**
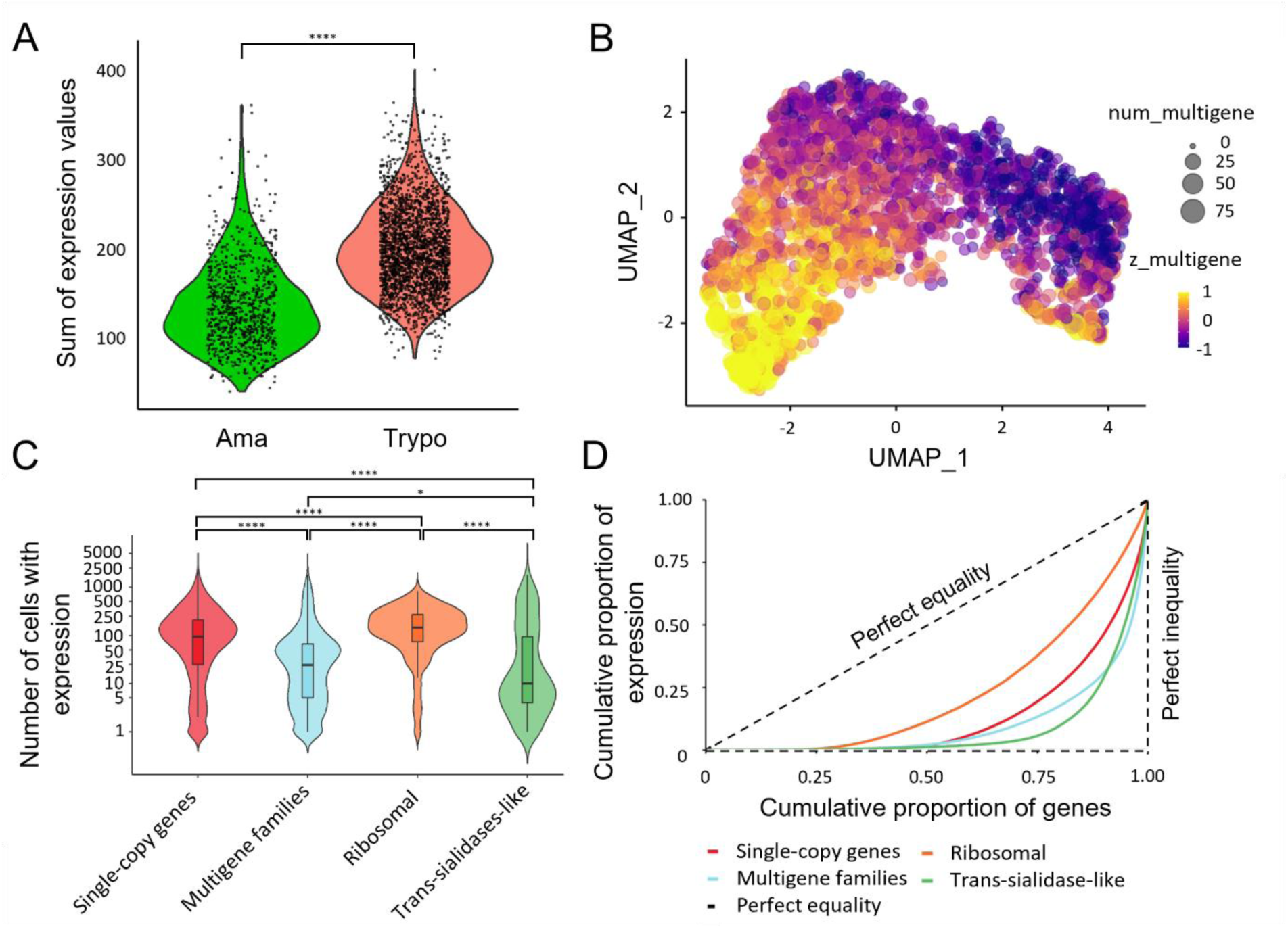
Overview of expression patterns across amastigote and trypomastigote cells. (a) Summatory of expression levels values from all multigene family genes for each cell from amastigote (Cluster 1) and trypomastigote (Cluster 0) cell populations (**** p < 0.0001, mean_Ama_ = 137.2, SD_Ama_ = 48.7, mean_Trypo_ = 201.1, SD_Trypo_ = 48.6, FC_Trypo/Ama_ = 1.5). (b) UMAP visualization of the expression patterns of multigene family genes; num_multigene indicates the number of multigene family genes detected per cell (genes with >0 UMI counts). Z_multigene reflects the relative expression level of multigene family genes per cell, calculated as the z-score-standardized sum of their UMI counts, such that positive values reflect above-average multigene family expression and negative values reflect below-average levels. (c) Violin plots showing the number of cells expressing a specific gene belonging to each group of genes: subsampled single-copy and multigene families, ribosomal genes and trans-sialidases. To avoid biases against size differences between single-copy and multigene family genes we generated a subsampled single-copy genes list, randomly selecting an equal number of genes as those from the multigene family’s gene set. The expression distribution of the subsampled single-copy genes is similar to the distribution of the entire dataset (* p < 0.05, **** p < 0.0001. See **Supplementary Table 2**). (d) Lorenz curves showing the cumulative proportion of gene expression relative to the cumulative proportion of genes for subsampled single-copy, multigene family genes, ribosomal protein coding genes and trans-sialidase-like genes. Genes were ordered by total expression, and the dashed line indicates perfect equality. Curves that deviate further from the diagonal reflect greater inequality, meaning that fewer genes account for most of the expression within each category. Statistically significant differences between groups for c) and d) are shown in **Supplementary Table 2**.

Within trypomastigotes, heterogenous expression of surface protein-coding gene families was observed, with high variation (compared to single-copy genes) in the number of cells in which each surface protein-coding gene was detected, as well as the total expression level in each cell (**Figure 2b and 2c**). Even though expression heterogeneity is also observed for single-copy genes (**Supplementary Figure 1a and Figure 1b**), probably due to the sampling biases that cause gene dropout in 10X Genomics technology, we investigated whether this phenomenon was more pronounced in surface multigene families. Therefore, we analyzed differences among single-copy and multigene family genes (together or grouped by multigene family) in trypomastigotes, in terms of the number of cells expressing each of the individual genes of each group (**Figure 2c, Supplementary Figure 1c and Supplementary Table 2**). In addition, we assessed expression inequality using Lorenz curves (**Figure 2d, Supplementary Figure 1d and Supplementary Table 2**) to evaluate how unevenly gene expression is distributed within each gene group. Ribosomal protein–coding genes were included as a control group.

Interestingly, compared to single-copy genes, especially ribosomal genes, multigene family genes showed a greater dispersion regarding the number of cells in which each gene was detected (**Figure 2c, Supplementary Figure 1c and Supplementary Table 2**), as well as a more pronounced deviation from the diagonal in the Lorenz curves (**Figure 2d, Supplementary Figure 1d)**, which represents perfect equality (reflected by different Gini indexes, see **Supplementary Table 2)**. Both observations indicate a higher expression heterogeneity for surface protein expression in the trypomastigote population. When multigene families were analyzed separately, several showed significant differences when compared to single-copy genes. Heterogeneity of the MASP family has already been suggested by studying clonal populations supporting our single cell results (31). Also, TcS genes exhibited a pronounced expression heterogeneity (**Supplementary Figures 1c and 1d**). TcS constitute a well-established and biologically central gene family in *T. cruzi*, playing key roles in host-parasite interactions (11,12,17). In this context, we next focused on a more detailed characterization of TcS expression heterogeneity in trypomastigotes.

### Expression pattern of the TcS superfamily

When we re-clustered the trypomastigote population based solely on TcS gene expression, we identified two sub-populations: trypomastigote cluster 0 (“Trypo_0” composed of 1186 cells), which over-expressed these genes compared to trypomastigote cluster 1 (“Trypo_1” composed of 1015 cells) (**Figure 3a**).

**Figure 3.**
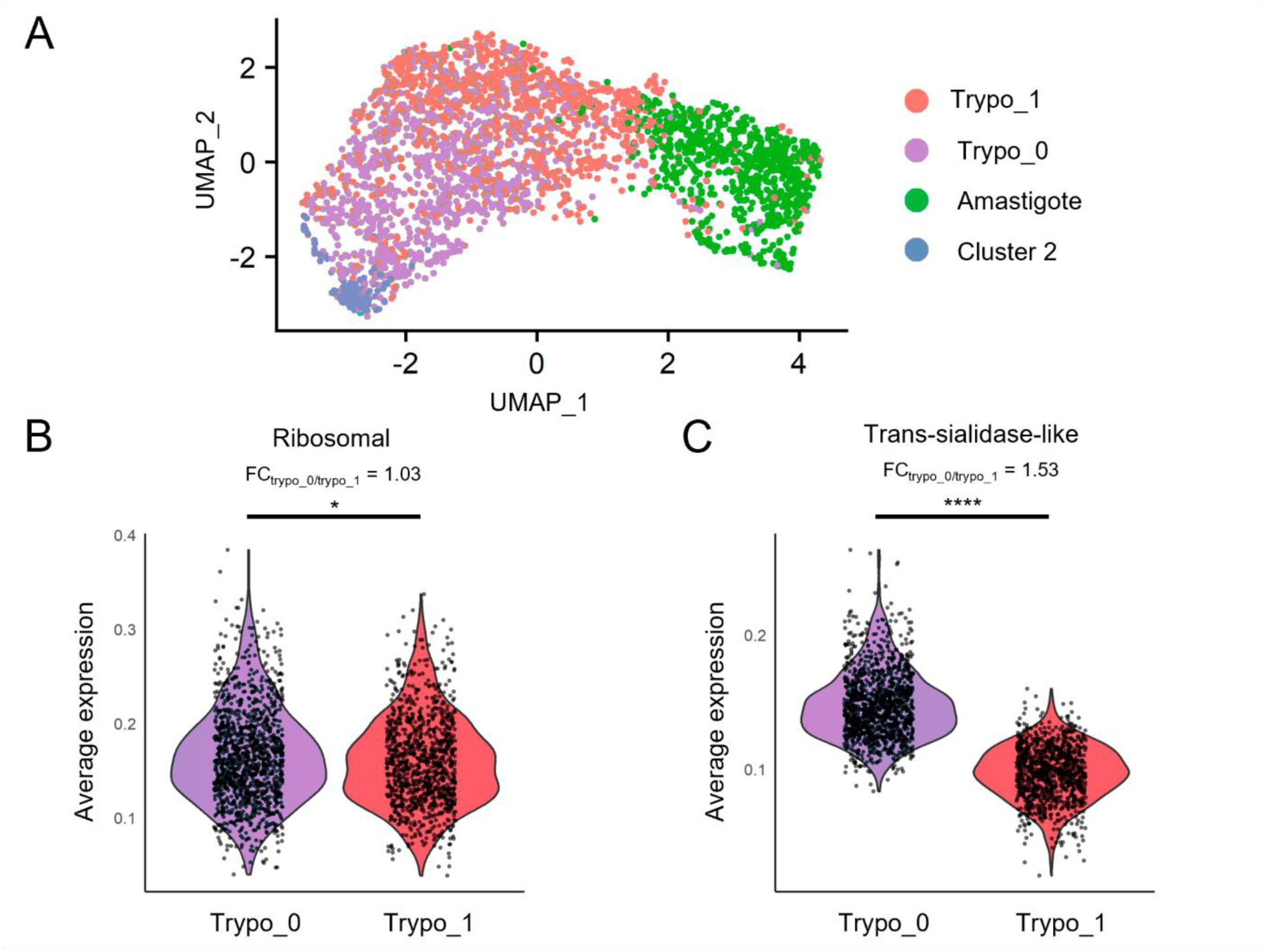
Trypomastigote sub-populations identified based on trans-sialidase expression profiles. (a) UMAP visualization colored by detected clusters based on gene expression profiles, with trypomastigote subpopulations identified. (b) Violin plot displaying average expression levels of ribosomal protein-coding genes across sub-populations. (c) Violin plot showing combined trans-sialidase expression levels for each sub-population. * p < 0.05, **** p < 0.0001.

The two trypomastigote sub-populations segregated by TcS expression show only slight differences in the expression of other gene categories, such as ribosomal protein-coding genes (FC = 1.03), transporters (FC = 1.09), polymerase-related genes (FC = 1.15), and phosphatases (FC = 1.10) (**Figure 3b and Supplementary Figure 2**, foldchange values correspond to Trypo_0/Trypo_1 expression ratios). Although there is a tendency for surface protein genes to be more expressed in cluster Trypo_0 (FC = 1.38), trans-sialidases displayed the highest fold change between the two trypomastigote subpopulations (FC = 1.53) (**Figure 3c and Supplementary Figure 2**). It is tempting to speculate that this may reflect different infectivity amongst trypomastigote subpopulations, consistent with reports of "broad" and "slender" forms (35). Recently, while this manuscript was being prepared, a similar approach by Laidlaw et al. described the phenomenon by clustering trypomastigote cells based on the expression of all genes (34). When applying this strategy to our data, their observation was reproduced.

When all cells are considered, most TcS genes are expressed, but in each individual cell only approximately 40 TcS genes were detected. Interestingly, we observed that both trypomastigote subpopulations contain a subgroup of TcS genes that are detected in a large portion of cells (>40%) (**Figure 4a, and Supplementary Table 3**), indicating high-level expression at the population level. This subset comprises 31 TcS genes belonging to subfamilies II-VI, and VIII (**Supplementary Table 3**). Consistent with our findings, Laidlaw et al. (34) reported a similar phenomenon, detecting on average approximately 40 TcS genes per cell, with a subset that are frequently detected across the trypomastigote population, further supporting the reproducibility of scRNA-seq results across studies. Gene dropouts in scRNA-seq experiments can generate apparently stochastic detection patterns for multigene family members, particularly affecting low-abundance transcripts, and thus represent an important potential confounder in the analysis of multigene families. To evaluate whether this technical effect could account for the observed TcS detection patterns, we examined the relationship between detection frequency and expression level. If random sampling were the primary driver, genes with similar average expression levels would be expected to exhibit comparable detection frequencies. However, we observed that many TcS genes with similar average expression are detected in markedly different proportions of cells (**Figure 4b**), arguing against a purely stochastic dropout model. Notably, among the 50 TcS genes with the highest average expression across cells, 62% are detected in fewer than 5% of cells (**Figure 4b**).

**Figure 4.**
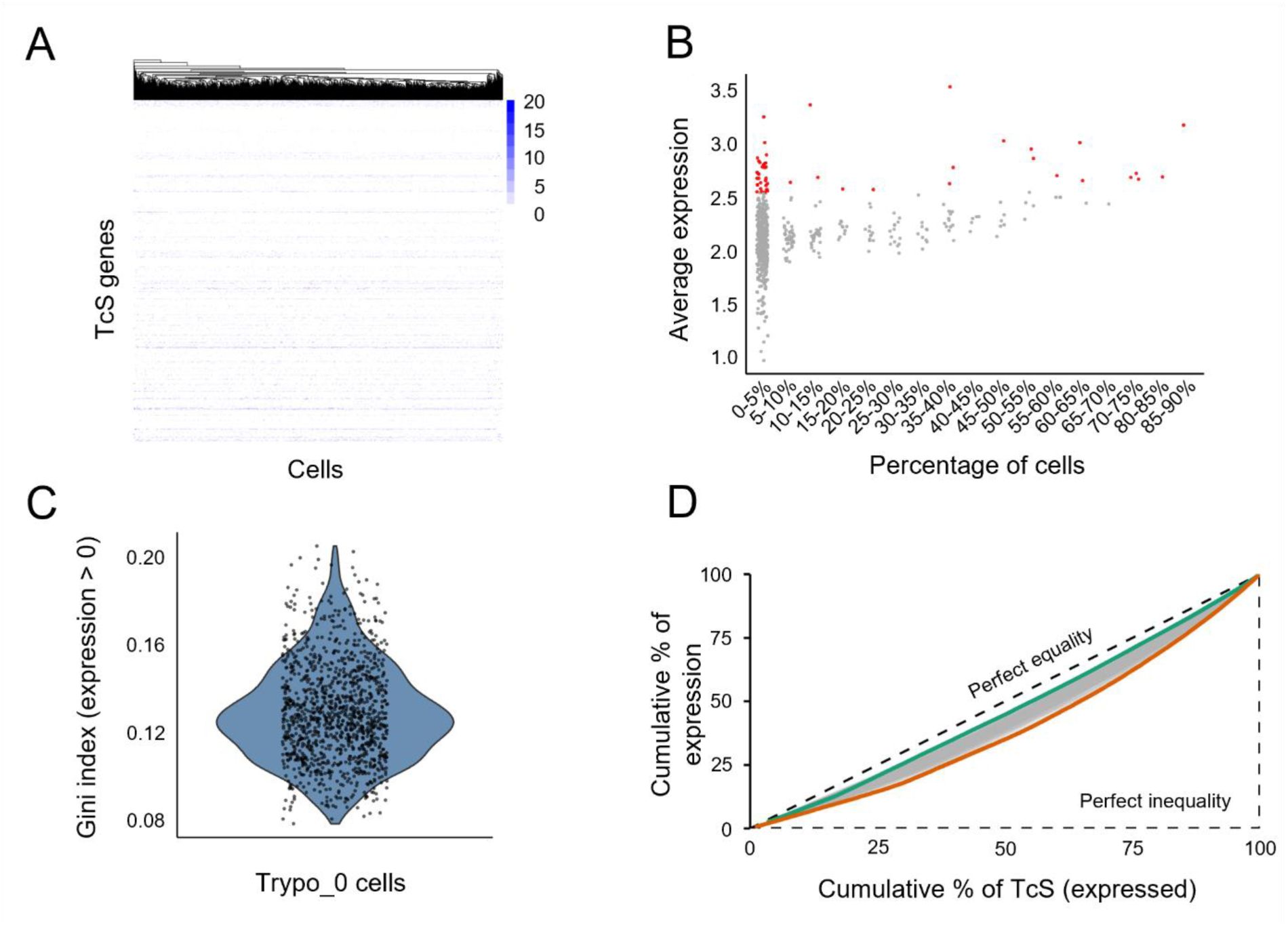
Overview of TcS gene expression patterns in Trypo_0 cells. (a) Heatmap displaying the expression of TcS genes in each cell that together account for 75% of total TcS gene expression within cluster Trypo_0. Cells are clustered by TcS expression profiles, with colors representing each gene’s percentage contribution to the cell’s total TcS expression. (b) Average expression of TcS genes grouped by the percentage of cells expressing each gene. In red are highlighted the top 50 TcS with highest average expression. (c) Gini index distribution for trypomastigotes cluster 0 (Trypo_0) cells considering only TcS detected in each cell. (d) Lorenz curves showing, for each cell in cluster Trypo_0, the cumulative proportion of total TcS expression as a function of the cumulative proportion of detected TcS genes. Genes were ordered by total expression, and the dashed line indicates perfect equality (i.e., all detected TcS genes contribute equally to the total TcS expression of a given cell). Green and orange curves correspond to cells with higher and lower expression equality, respectively.

At the single-cell level, we found that detected TcS genes contribute relatively evenly to the total family expression within each cell, as indicated by the low average Gini index of TcS expression levels across each cell (**Figures 4c and 4d**). This suggests that, within each cell, total TcS expression is shared across the detected TcS genes rather than being dominated by a few very highly expressed transcripts in that cell. Consistent with this interpretation, most TcS genes detected in a high fraction of cells in our scRNA-seq dataset also rank among the most highly expressed TcS genes in an independent bulk RNA-seq study (**Supplementary Figure 3a**). This highlights a key limitation of bulk RNA-seq, as it may wrongly indicate that a set of few genes are highly expressed in each cell, while in fact, TcS expression is evenly distributed among all detected family members, regardless of the number of cells expressing each gene. Taken together with the results in **Figure 4b**, this observation further supports the conclusion that the observed expression pattern is unlikely to arise from a purely stochastic detection of TcS mRNAs.

While our data indicates transcriptional heterogeneity, it is important to note that future studies will be required to determine the extent to which this diversity translates to protein expression on the parasite surface. Consistent with our observations, recent proteomic analyses (36) report pronounced variability in the expression of surface proteins, including TcS, suggesting that the heterogeneous transcriptional patterns observed here likely reflect biologically relevant differences.

When analyzing each trypomastigote subpopulation, no coordinated expression among specific TcS members was observed, as no subclusters of cells were identified based on TcS detection profiles. Even when clustering was restricted to genes detected in more than 40% of cells, no clear subclusters of cells were identified (**Supplementary Figure 3b**).

As discussed in the Introduction, although post-transcriptional control remains a central mechanism of gene expression regulation in *T. cruzi*, increasing evidence in recent years has revealed an important contribution of epigenetic mechanisms that modulate gene expression at the transcriptional level. (4–8). Specifically, the loci for TcS genes and other gene families of surface proteins are mostly grouped in specific genomic compartments (9) that are regulated epigenetically by the activation or silencing of chromatin folding domains (4). The TcS that are detected in a high percentage of cells are mostly dispersed throughout the genome (**Supplementary Table 3**). This suggests that their preferential expression is likely not due to colocalization in one or a few ubiquitous activated chromatin-folding domains. Nevertheless, mapping the genomic locations of TcS genes detected in a high proportion of cells revealed that most are flanked by core genes (**Figure 5**). The core compartment is enriched in conserved, single-copy genes that typically show more constitutive expression (9) and, as observed in this study, lower cell-to-cell variability. In contrast, TcS genes that are detected less frequently are preferentially located in the disruptive compartment (**Figure 5**), which is enriched in lineage-specific multigene families and associated with more variable, stage-specific, and potentially stochastic expression under tighter epigenetic control (4,9,18,36). Together, these findings suggest that the higher cellular prevalence of certain TcS transcripts is unlikely to be driven by colocalization within a small number of ubiquitously active chromatin domains but may instead reflect distinct regulatory regimes between the core and disruptive compartments. Future studies integrating single-cell chromatin profiling with scRNA-seq will be required to directly test this model.

**Figure 5.**
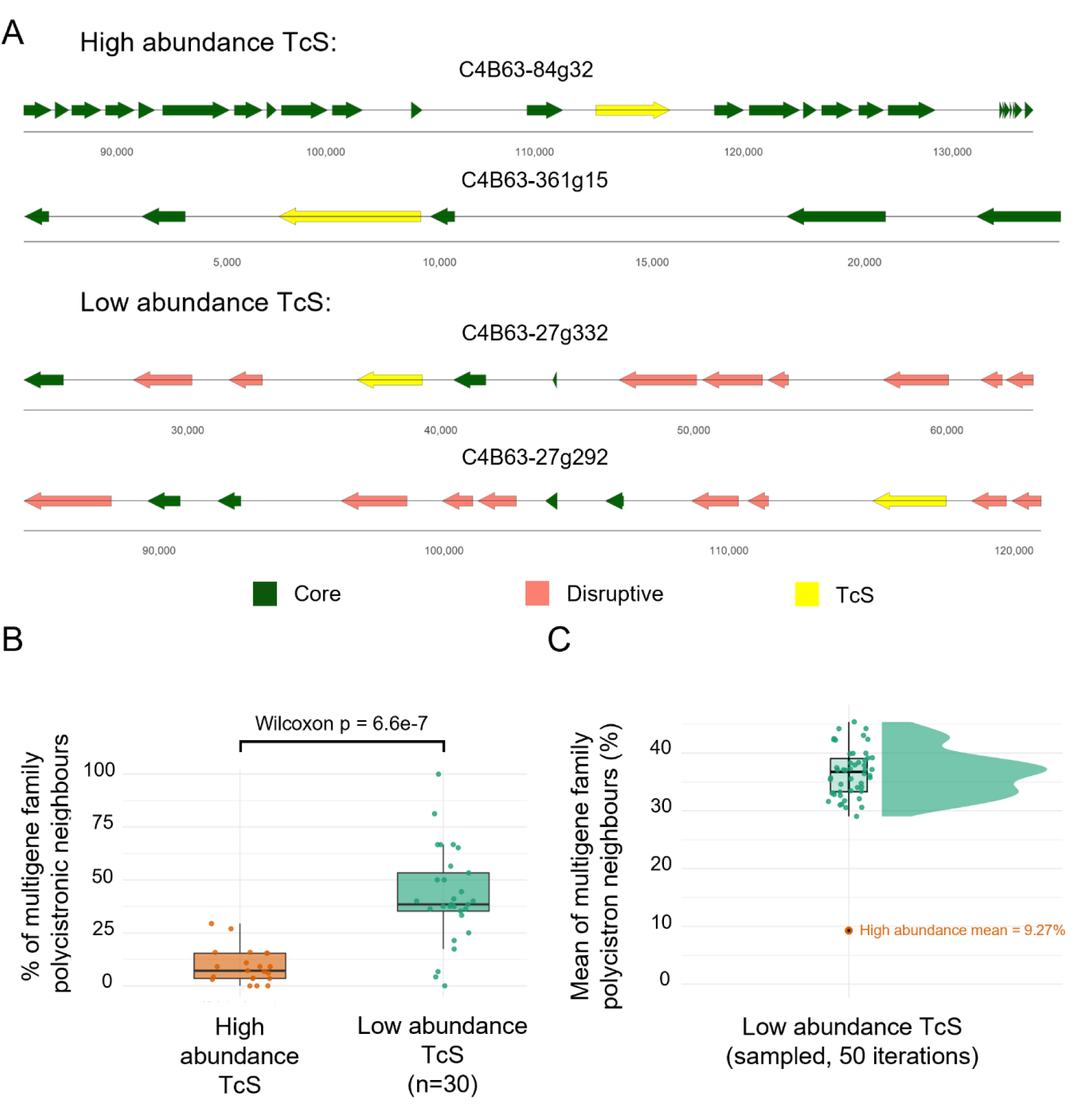
Genomic context and neighborhood composition of frequently detected versus lowly detected TcS loci. (a) Representative genomic loci of frequently detected (top, high abundance) and lowly detected (bottom, low abundance) TcS genes. Genes are shown as arrows, colored according to genomic compartment: core (dark green), disruptive (salmon), and TcS genes under analysis (yellow). Chromosomal coordinates are indicated below each locus. (b) Comparison of the percentage of multigene-family neighbours within polycistronic transcription units containing frequently detected and lowly detected TcS genes. Lowly detected TcS genes were subsampled to *n* = 30. Wilcoxon rank-sum test: *p* = 6.6 × 10⁻⁷. (c) Mean percentage of multigene-family neighbours in polycistrons calculated from 50 random subsets (*n* = 30) of lowly detected TcS genes (mean = 37.76%). The orange dot indicates the corresponding mean for frequently detected TcS genes (9.27%).

### Final remarks

The expression of surface protein–coding genes varied across parasite developmental stages. In particular, genes belonging to the TcS superfamily, which play key roles in host–parasite interactions, exhibited marked heterogeneity in expression among trypomastigotes. Notably, while most TcS genes were detected only in a small fraction of cells, a limited subset was frequently detected across the population. This pattern indicates that TcS expression is not uniform, suggesting the existence of distinct regulatory regimes within the family. Consistent with previous single-cell and bulk RNA-seq studies (34,36), the lack of coordinated expression among TcS members points to a complex regulatory framework that may enable functional diversification among *T. cruzi* subpopulations. Importantly, our analyses indicate that the frequent detection of this TcS subset may be partially explained by their preferential localization within the core genomic compartment, which is associated with more permissive transcriptional environments.

Taken together, our results demonstrate the sensitivity of scRNA-seq for resolving parasite life stages and their associated transcriptional programs, while also pointing to complex regulation of surface protein expression, particularly within the TcS family. Although these results should be considered with caution, as technical limitations inherent to scRNA-seq may influence the observed expression patterns, our interpretation is consistent with all observations presented here and aligns with emerging evidence from independent studies. Collectively, this supports a working hypothesis in which heterogeneous TcS expression may contribute to immune evasion and pathogenicity through a bet-hedging strategy.

## Materials and Methods

### Trypanosoma cruzi and mammalian cell culture

Epimastigote forms of *Trypanosoma cruzi* strain Dm28c were derived from axenic cultures cultivated in Brain-Heart Infusion medium (BHI, Oxoid) supplemented with 10% heat-inactivated fetal bovine serum (FBS, Capricorn), penicillin (100 units/mL) and streptomycin (100 µg/mL) as described (37). Cultures were diluted 1/10 with fresh BHI medium every 3 days and maintained at 28°C.

Myoblast rat cell line H9c2 (ATCC CRL-1446) was maintained in hgDMEM medium (Gibco) supplemented with 10% heat-inactivated fetal bovine serum, penicillin (100 units/mL) and streptomycin (100 μg/mL) at 37°C in a humidified 5% CO_2_ incubator. Confluent cells were washed with 1X phosphate-buffered saline (1X PBS), incubated for 5 min with trypsin-EDTA (Gibco), diluted with culture medium and re-plated for maintenance.

Mycoplasma contamination in cell lines was regularly monitored using MycoAlert® Mycoplasma Detection Kit (Lonza), following the manufacturer’s protocol.

### Isolation and purification of cellular trypomastigotes and intracellular amastigotes

Late stationary phase epimastigotes of Dm28c strain were used to infect H9c2 cells for a primary infection. Six days post-infection, cellular trypomastigotes were obtained from the supernatant and were used to infect 50% confluent H9c2 cells at a 10:1 rate. Twenty-four hours post-infection the cell culture was washed twice with PBS 1X to remove any remaining extracellular parasites and maintained with fresh hgDMEM at 37°C in a humidified 5% CO_2_ incubator.

For amastigote purification, infected H9c2 cells incubated for 48 hours post infection were washed with 1X PBS and incubated with trypsin-EDTA for 5 minutes at 37°C. The trypsinization was stopped by adding an equal volume of hgDMEM with 10% FBS. The cell suspension was repeatedly passed through a 27-gauge needle attached to a 30-mL syringe until complete cell disruption was confirmed under the microscope. The supernatant, containing free amastigotes, was collected and centrifuged at 500xg for 10 minutes at 4°C to remove large host-cell debris. The resulting supernatant was then centrifuged at 4,000xg for 10 minutes at 4°C, and the amastigote-containing pellet was washed twice in chilled 1X DPBS (Dulbecco’s Phosphate-Buffered Saline, No Calcium, No Magnesium) and resuspended in 1X DPBS at 200 μL per 1×10^6^ cells, ready for the fixation step.

Cellular trypomastigotes derived from infected H9c2 cells and present in the cell supernatant fraction were collected and centrifuged at 500xg for 10 minutes at 4°C to remove large host-cell debris. The washing and resuspension steps in DPBS were performed as previously described for amastigotes.

### scRNA-seq library preparation and sequencing

Cell fixation was performed using the Methanol Fixation Protocol for Single-Cell RNA Sequencing (38), after resuspension in DPBS as described above, as recommended by 10X Genomics technical support. Briefly, chilled 100% Methanol (for HPLC, ≥99.9%, Millipore) was added drop by drop (1×10^6^ cells in 800 μl) and incubated at -20°C for 30 min. For rehydration, fixed cells were first equilibrated at 4°C and then centrifuged at 4,000xg for 5 min at 4°C, the supernatant was discarded and Wash-Resuspension Buffer (3X SSC in Nuclease-free Water, 0.04% UltraPure Bovine Serum Albumin, 1mM DTT, and 0.2 U/ml RNase Inhibitor) was added to the pellet. Cell debris and large clumps were eliminated by passing the sample through a 40 μm Flowmi Cell Strainer.

10X Genomics library preparation was performed at the service provided by the Instituto de Biología y Medicina Experimental (IBYME, Argentina). The library was sequenced by a service provider (Macrogen, Korea) in a HiSeq2500 equipment (two lanes), generating approximately880 million reads of 91 bp. Raw data is available in the SRA (https://www.ncbi.nlm.nih.gov/sra/) under BioProject PRJNA1200704.

### Transcript quantification

*T. cruzi* 2018 Dm28c genome (9) (release 62, TriTrypDB (39)) and *T. cruzi* Dm28c maxi circle kDNA sequence (40) were combined and used as reference. To improve the proportion of reads assigned to genes, 11,362 3’UTR regions of the coding sequences (CDS) were annotated using peaks2UTR (41). Gene expression quantification was performed by pseudoalignment using kallisto bustools (42), with the options --filter to remove potential noise from environmental RNA and --em to apply the Expectation-Maximization (EM) algorithm. Default values were used for the rest of the parameters. This algorithm outperforms other mapping software in handling multimapping reads, enabling more accurate quantification of multigene families (42). This resulted in 321 million reads mapped to the transcriptome.

### scRNA-seq data processing and analysis

Count matrices from two technical replicates obtained by sequencing the same library were merged: the common barcodes across both datasets were retained, and the count matrices were combined to generate a unified dataset. Subsequently, the following metrics were calculated: nUMI, nGene, and mitoRatio, as well as the log10 of genes per UMI (log10GenesperUMI). A filtering criterion was applied, retaining cells with nUMI > 1200, nGene > 100, log10GenesperUMI > 0.8, and mitoRatio < 0.1. Ribosomal rRNA genes were excluded from subsequent analyses. Data normalization and scaling were performed using the Seurat R version 5 package (43), employing NormalizeData and ScaleData functions.

After quality filtering, 3,192 single-cell transcriptomes were retained with an average of 8004 reads mapped per cell. In total, 14,321 genes were detected across all cells (93.5% of the 15,319 annotated protein-coding genes in the *T. cruzi* 2018 Dm28c genome). Per cell, we observed a mean of 1,088 detected genes and 2,461 UMIs, which corresponds to ∼7.1% of the annotated protein-coding gene.

FindNeighbors function was used to construct a k-nearest neighbours graph of cells using 10 principal components (PCs) and Louvain algorithm was employed for cell clustering using FindClusters function. Additionally, doublets were identified and removed using the DoubletFinder R package (44); only 41 doublets were detected, all belonging to cluster 2.

To define marker genes, the FindAllMarkers function from Seurat package was used, selecting those with an adjusted p-value < 0.05 and a log2 fold change (log2FC) > 1. To validate these markers, bulk RNA-seq data of Dm28c was incorporated (NCBI BioProject ID PRJNA850400 [24]). Transcripts were quantified using Kallisto (45) (with -b 100 option to perform 100 bootstraps), followed by differential expression analysis conducted with Sleuth (46). Genes with an adjusted p-value < 0.05 and |log2FC| > 0.25 were filtered for further analysis. Finally, the gene IDs of the Seurat-defined markers were cross-referenced with the IDs of the differentially expressed genes obtained from the bulk RNA-seq analysis to corroborate stage-specific gene expression. For 2D visualization of cell clusters and gene markers expression profiles across cells, UMAP projection was employed (47).

Gene IDs corresponding to multigene families (TcS, MASP, Mucins, GP63, RHS and DGF) were obtained by text searches using the current genome annotation on the “description” field. Throughout the manuscript, these genes are referred to as either surface protein genes or multigene family genes. Single and low copy number genes were defined as those that did not belong to the latter gene list. For simplicity “single-copy” will be used to refer to these genes throughout the manuscript. To avoid biases arising from differences in list size between single-copy and multigene family gene sets, we generated a subsampled single-copy gene list by randomly selecting the same number of genes as in the multigene family set. The expression distribution of this subsampled single-copy gene set is similar to that of the full dataset.

Data processing was conducted using R version 4.2.0. Statistical analyses were performed using the Wilcoxon rank-sum test and p-values < 0.05 were considered statistically significant.

To study expression inequality, we plotted Lorenz curves (48) and applied the Gini index, a metric originally developed in the field of economics (49). Both metrics serve as a measure of expression heterogeneity, and we employed it in two distinct contexts: first, to evaluate the degree of inequality in the distribution of a gene’s expression levels across individual cells (**Figure 2d, Supplementary Figure 1d and Supplementary Table 2**); and second, to assess the extent to which a given cell expresses individual genes at varying rates (**Figure 4c**).

TcS genomic localization analysis was performed by defining directional gene clusters from the GFF annotation: protein-coding genes were ordered by coordinate per contig and consecutive genes on the same strand were grouped into a DGC, with a strand-switch (change from “+” to “-” or vice versa) marking DGC boundaries. For each gene, we calculated the percentage of its polycistronic neighbours (genes within the same DGC) that belong to the multigene family gene set previously defined and compared high-abundance TcS vs low-abundance TcS (sampling n = 30). For statistical comparisons, we assessed the robustness of the analysis by sampling 50 iterations of n = 30 low-abudance TcS and distributions were compared to the high-abundance set using two-sided Wilcoxon rank-sum tests with Benjamini-Hochberg correction.

## Author Contributions

LI: Data Curation, Formal Analysis, Investigation, Methodology, Visualization, Writing-Original Draft Preparation, Writing-Review & Editing; LB: Methodology, Investigation, Writing-Review & Editing; VAC: Methodology, Investigation, Writing-Review & Editing; JG: Methodology, Formal Analysis, Writing-Review & Editing; GR: Methodology, Writing-Review & Editing; VMH: Methodology, Writing-Review & Editing; MAD: Methodology, Resources, Writing-Review & Editing; JSS: Methodology, Resources, Writing-Review & Editing; JDG: Conceptualization, Methodology, Visualization, Writing-Original Draft Preparation, Writing-Review & Editing; PS: Conceptualization, Funding Acquisition, Methodology, Formal Analysis, Project Administration, Resources, Supervision, Validation, Writing-Original Draft Preparation, Writing-Review & Editing

## Supplementary Figures

**Supplementary Figure 1.**
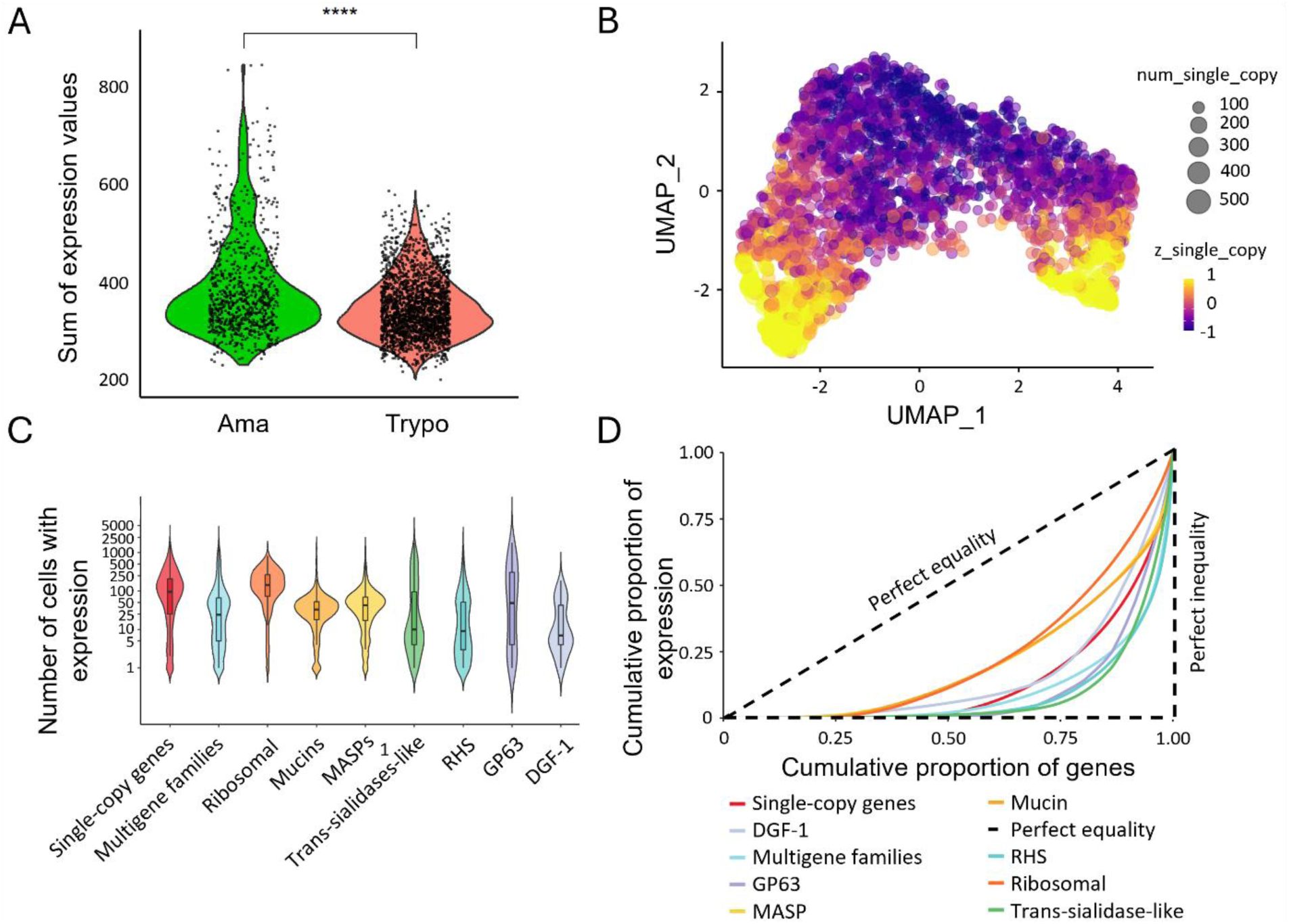
(a) Summatory of expression levels values from subsampled single-copy genes for each cell from amastigote (Cluster 1) and trypomastigote (Cluster 0) cell populations (**** p < 0.0001, meanAma = 394.9, SDAma = 101.7, meanTrypo = 353.1, SDTrypo = 62.1, FCAma/Trypo = 1.1). (b) UMAP projection for 2D visualization of core gene expression among cells, ; num_multigene indicates the number of multigene family genes detected in each cell, whereas z_multigene indicates the expression levels calculated by summing the UMI counts of all multigene family genes in each cell and then standardizing this value using a z-score transformation, such that positive values reflect above-average multigene family expression and negative values reflect below-average levels. (c) Violin plots showing the number of cells expressing a specific gene belonging to each group of genes: subsampled single-copy and multigene families, ribosomal genes and different multigene families. (d) Lorenz curves showing the cumulative proportion of gene expression relative to the cumulative proportion of genes for subsampled single-copy, multigene family genes (together or grouped by multigene family) and ribosomal protein coding genes. Genes were ordered by total expression, and the dashed line indicates perfect equality. Curves that deviate further from the diagonal reflect greater inequality, meaning that fewer genes account for most of the expression within each category. Statistically significant differences between groups for c) and d) are shown in **Supplementary Table 2**.

**Supplementary Figure 2.**
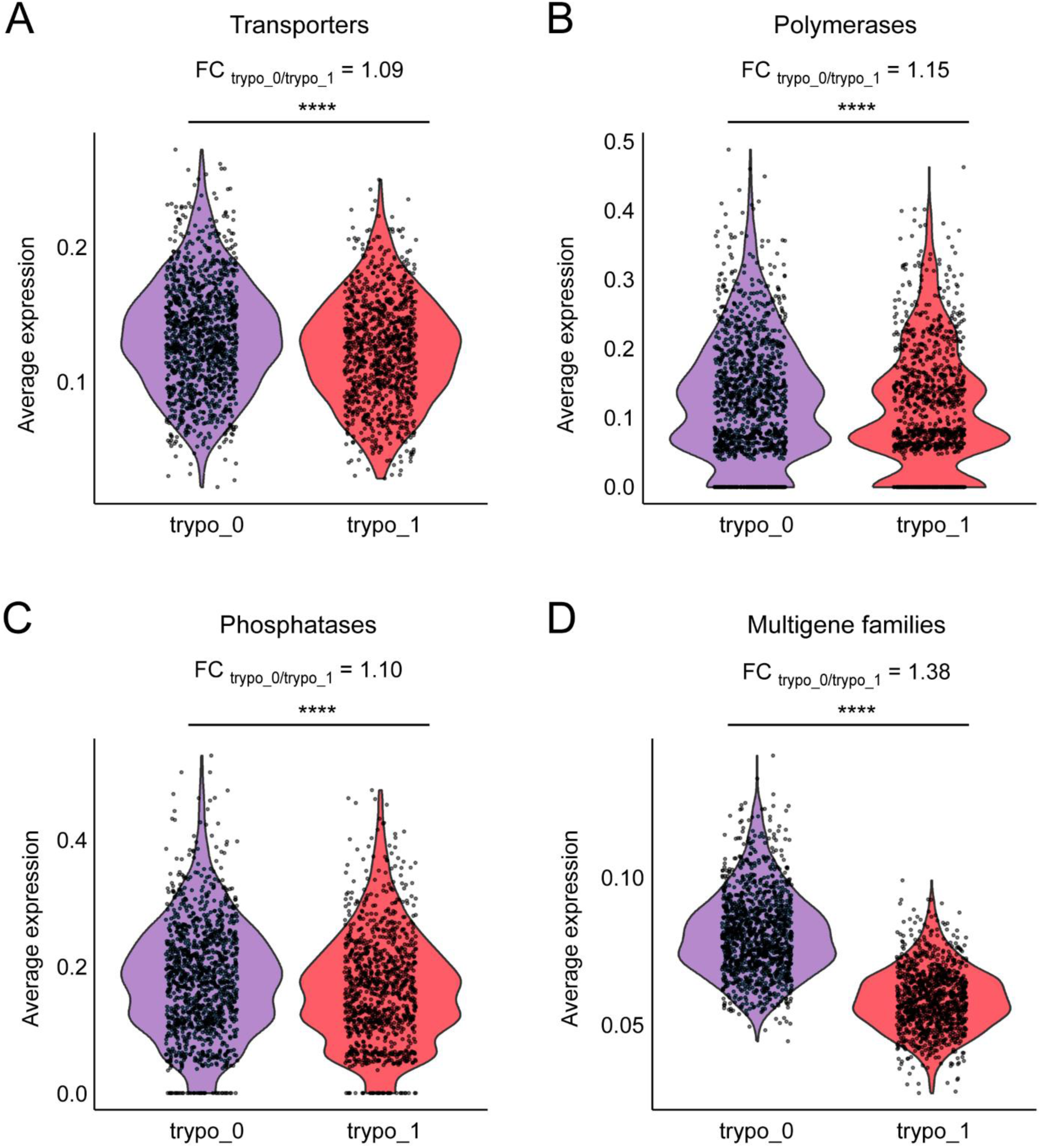
Trypomastigote sub-clusters identified based on trans-sialidase expression profiles. Violin plots displaying average expression levels across sub-clusters and associated fold changes (FC_trypo_0/trypo_1_) of (a) transporters coding genes, (b) DNA and RNA polymerase-associated protein coding genes, (c) phosphatases coding genes and (d) multigene family genes. **** p < 0.0001.

**Supplementary Figure 3.**
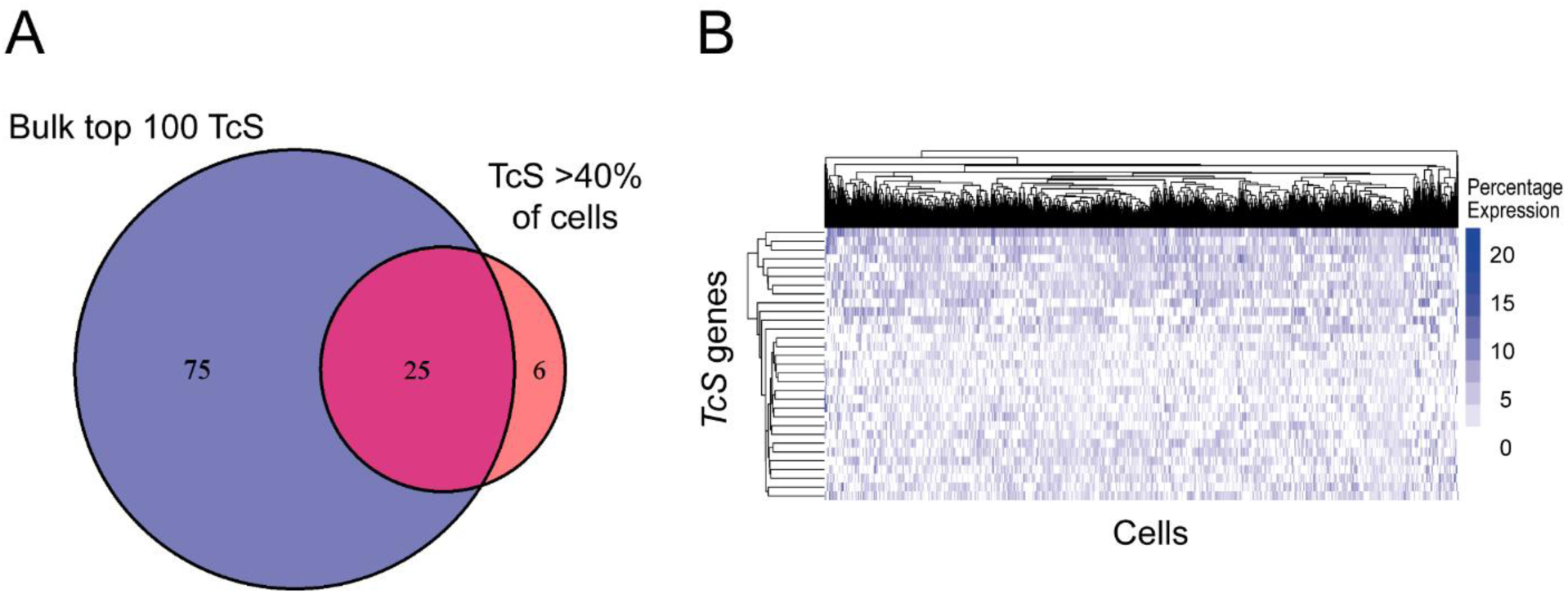
(a) Venn diagram showing the overlap between the top 100 most expressed TcS from bulk RNA-seq data and TcS expressed in more than 40% of cells from cluster trypo_0, and (b) Heatmap displaying the expression of TcS genes in each cell that together account for 75% of total TcS gene expression and are expressed in more than 40% of cells within cluster trypo_0. Cells are clustered by TcS expression profiles, with colors representing each gene’s percentage contribution to the cell’s expression.

